# Timing of metabolomics-driven supplementation strategies affects protein expression in *E. coli-*based cell-free expression systems

**DOI:** 10.64898/2026.08.26.746766

**Authors:** Soor R. Vora, Mark P. Styczynski

## Abstract

While *in vivo* synthesis of biologic therapeutics has been broadly successful, it is limited by biological constraints of the cells and by the complexity, time, and cost of implementing the pipeline from discovery through manufacturing. Cell-free expression systems (CFES), which use cellular transcription and translation machinery to express proteins *in vitro*, offer a promising alternative approach that could improve robustness and modularity in that pipeline. However, current benchmark CFES productivity is well below the theoretical capacity of the input nucleotides and amino acids. Efforts to address this issue are hindered by limited understanding of the extent of enzymatic activity in CFES beyond gene expression, as previous work has shown that metabolic enzymes in cell-free lysates cause substantial background metabolic activity that influences protein expression. Here, we hypothesized that the inflection point of protein expression is a critical timescale for CFES metabolism. We performed metabolomics characterization of CFES reactions, finding significant metabolic changes at the inflection point. Driven by these findings, we sought to identify supplements that could be added to the cell-free reaction to avoid metabolic limitations. We found that amino acid supplementation increased expression productivity and lifetime only when added after the inflection point, and actually hurt expression when added before the inflection point. We found similar supplementation timing impacts for some other metabolites as well. These findings show that endogenous metabolism and supplementation timing are deeply interconnected and are critical considerations in CFES optimization, and that metabolomics-informed fed-batch supplementation is a potentially valuable strategy to improve reaction productivity.

**Graphical Abstract:** 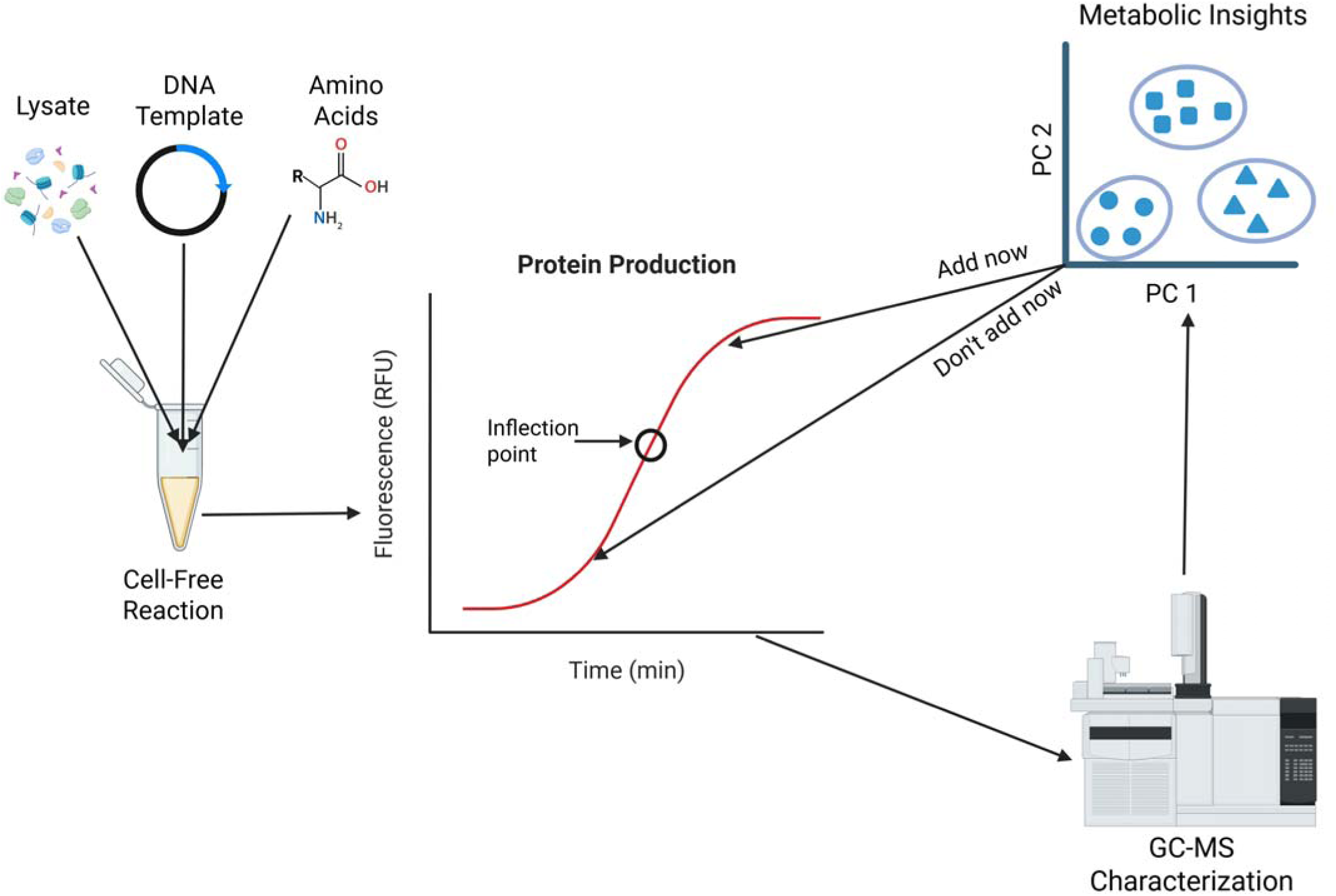

## Introduction

Biosynthesis of valuable molecules has many impactful applications, including the manufacturing of biologic drugs that are increasingly dominating the therapeutic landscape.^1^ *In vivo* production of biologics using whole cells is a powerful approach but is limited by both the requirement that the cells must remain viable during production as well as the extensive cell-line engineering requirements to program whole-cell biocatalysts to synthesize different proteins.^2^

Specifically, producing testable amounts of a candidate biologic requires a long and expensive process to create cell lines capable of making that new protein. Cell-free expression systems (CFES) are an emerging technology that provide an *in vitro* alternative to whole-cell biocatalysts. These systems use cellular extracts containing transcription and translation machinery to express proteins *in vitro*, with expression of a new protein possible merely by changing the DNA template provided to the reaction.^3^ *Escherichia coli (E. coli)-*based CFES are a popular choice due to their robustness, tunability, and low cost.^4,5^ The protein expression curve for these systems is approximately sigmoidal, with initially accelerating production due to accumulation of mRNA followed by a period of high protein productivity that eventually slows down and then plateaus.

While the robustness and modularity inherent to CFES have enabled their use in many applications and make them promising for producing testable or even manufacturing-scale quantities of biologics, CFES still face barriers to broad adoption. Most importantly, current benchmark titers for protein expression are well below theoretical maximum production based on stoichiometry of the amino acids and nucleotides provided as input to the reaction. Previous work has proposed and tested some potential causes of this limitation, including depletion of key metabolites and accumulation of toxic metabolites.^6,7^ Despite ongoing efforts at optimization, protein expression still plateaus well before CFES reach their theoretical protein production capacity, representing an inherent limit for all CFES-synthesized proteins that undercuts any product-specific metabolic engineering efforts that may be taken to enable higher titers of a specific protein.^8^

These efforts to improve cell-free system productivity are hindered by limited understanding of the extent of enzymatic activity in CFES beyond gene expression. The cellular transcription and translation machinery are present in cell-free lysates because those lysates retain the soluble protein fraction of the cell, which in turn contains metabolic enzymes from the cells. Previous work has demonstrated that these enzymes cause metabolic activity that influences cell-free expression, but the metabolic behavior and dynamics of these systems are only beginning to be characterized.^11,12^ Metabolomics, the systems-scale study of the small molecules (metabolites) in a biological system, is a powerful way to characterize that metabolic activity and identify potential connections with physiological outputs.^9,10^

In this work, we hypothesized that the inflection point of protein expression is a critical timescale for CFES metabolism that connects to the inability of CFES to reach their theoretical maximum production. We found that the metabolic activity of CFES reactions shifts significantly at the inflection point, suggesting the possibility of a metabolic trigger that leads to decreased expression activity. With that information, we proposed and implemented a metabolomics-informed fed-batch CFES reaction strategy, finding that controlled addition of metabolic resources can significantly improve CFES productivity and lifetime depending on the timing of that addition.

## Results

### Metabolic dynamics around the expression inflection point

We hypothesized that the inflection point of protein expression is a critical timescale for CFES metabolism and total protein production. Figure 1B shows the early portion of a representative cell-free protein production profile for a 210 μL cell-free reaction in a 96-well plate with sfGFP expression driven by a strong T7 promoter. The initial increase in expression rate occurs due to accumulation of mRNA transcripts leading to an acceleration of total expression. As transcription slows down but translation continues, the reaction enters an approximately linear regime. The expression rate then starts to slowly decrease until it eventually plateaus (not shown in Figure 1B). However, this decrease in expression rate occurs while there are still sufficient nucleotides and amino acids necessary for expression, signaling some other type of limitation.

**Figure 1:**
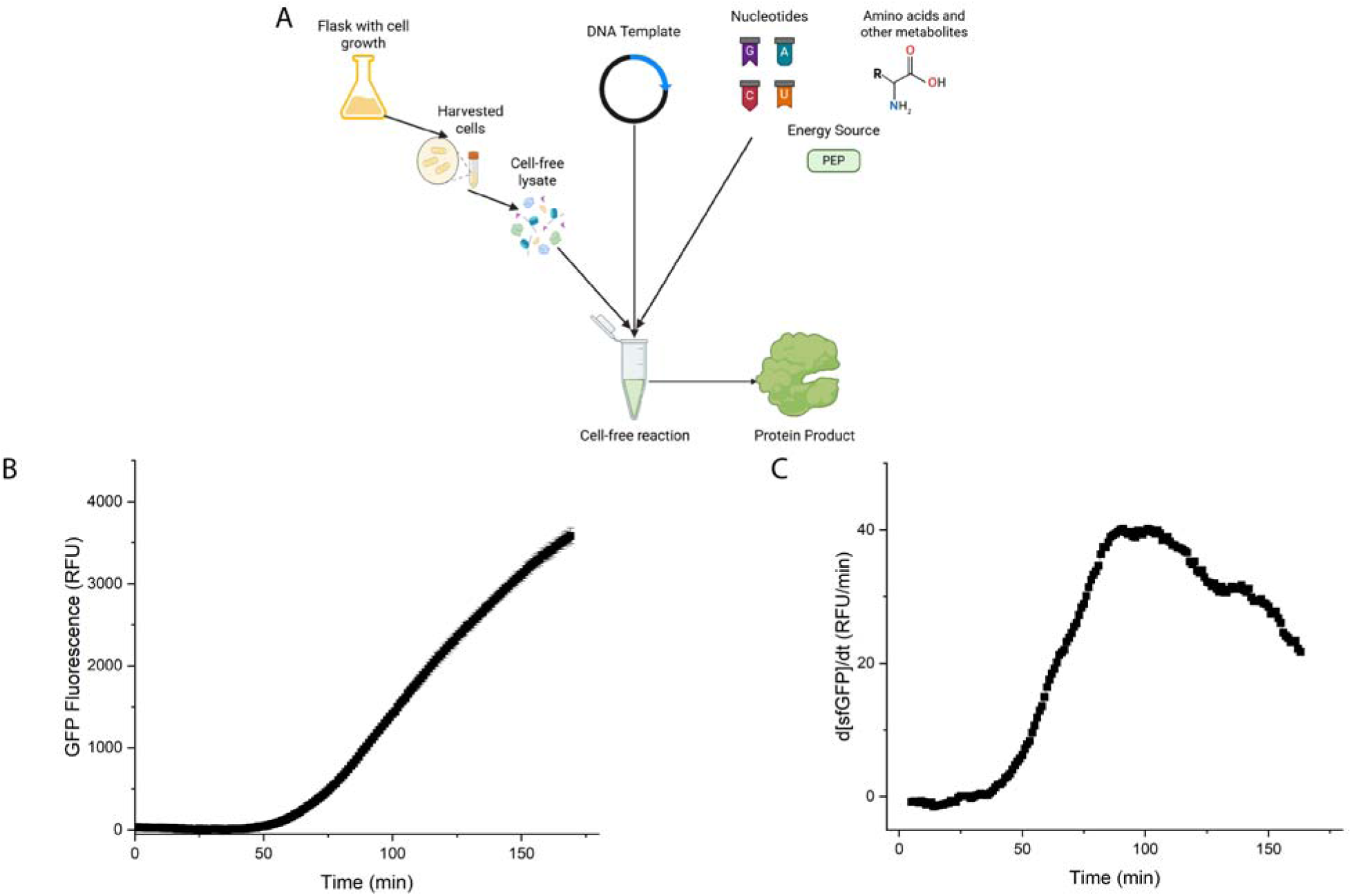
Schematic and expression dynamics of *E. coli*-based CFES in metabolomics-scale reaction volumes. A) Overview of reaction components for crude extract-based cell-free systems. Figure made in BioRender.^21^ B) Representative sfGFP fluorescence over time (error bars represent ± one standard deviation and C) rate of change of average fluorescence over time show an inflection point at approximately 100 minutes.

The inflection point of the expression profile in Figure 1b is clearly visible as a local maximum in the derivative of the fluorescence curve, which in this case occurs at approximately 100 minutes (Figure 1C). The inflection point varies depending on several conditions including reporter promoter strength, reaction scale, and reporter plasmid concentration (Table 1), as well as the operator running the reaction (data not shown). We selected 1 nM of T7-driven sfGFP as the condition we focused our analysis on as it is representative of a typical protein expression or prototyping condition, though we note that in our experience the metabolic profile is only weakly dependent on plasmid concentration, suggesting that similar results would be observed in other conditions.^11^ When characterizing reactions via metabolomics, we used a 210 μL reaction volume to ensure we had sufficient sample for GC-MS analyses. We focused most of our experiments on times around the inflection point (100 min when performing metabolomics analyses, 55 min when using smaller volumes to test only for expression).

**Table 1:** Time required to reach the inflection point of protein production for different plasmid concentrations, promoter strengths, and reaction volumes. The italicized inflection times are for the two conditions primarily used in this study. Numbers shown are averages of reaction triplicates paired with corresponding standard deviations.

| | 10 μL reaction inflection times (min) | | | 210 μL reaction inflection times (min) | | |
| --- | --- | --- | --- | --- | --- | --- |
| Plasmid<br>Concentration (nM) | Weak $\sigma^{70}$ | Strong $\sigma^{70}$ | Strong T7 | Weak $\sigma^{70}$ | Strong $\sigma^{70}$ | Strong T7 |
| 0.5 | 170.3±3.0 | 68.3±2.5 | 62.7±3.5 | 229.7±4.5 | 124.0±4.0 | 114.3±3.5 |
| 1 | 69.0±2.0 | 68.0±2.0 | *55.0±2.0* | 125.33±5.5 | 123.3±3.5 | *100.0±3.0* |
| 2.5 | 69.3±0.6 | 57.7±1.5 | 45.7±2.7 | 127±3.0 | 100.0±3.0 | 96±4.0 |
| 5 | 70.7±0.6 | 49.0±2.0 | 38.0±2.1 | 128.3±3.5 | 95.3±3.1 | 89.7±2.5 |
| 10 | 69.0±2.0 | 40.0±1.0 | 24.3±0.6 | 125.0±1.0 | 92.0±3.0 | 85.0±2.0 |

There is a clear shift in metabolic dynamics at the inflection point of cell-free protein production, as seen in Figure 2. After an initial period of substantial metabolic changes in the first 40 minutes of the reaction, the metabolome remains relatively constant over the next 60 minutes. However, right after the inflection point (at 100 minutes) there is an additional substantial change in metabolomic profiles. As seen in Figure 2A, the dimensionally reduced metabolic profile starts increasing in principal component 2 at the inflection point after having decreased from the beginning of the reaction. Similarly, in analyzing metabolite changes between consecutive time points for potential pathway enrichment, there is a spike in enriched pathways immediately after the inflection point (Figure 2B).

**Figure 2.**
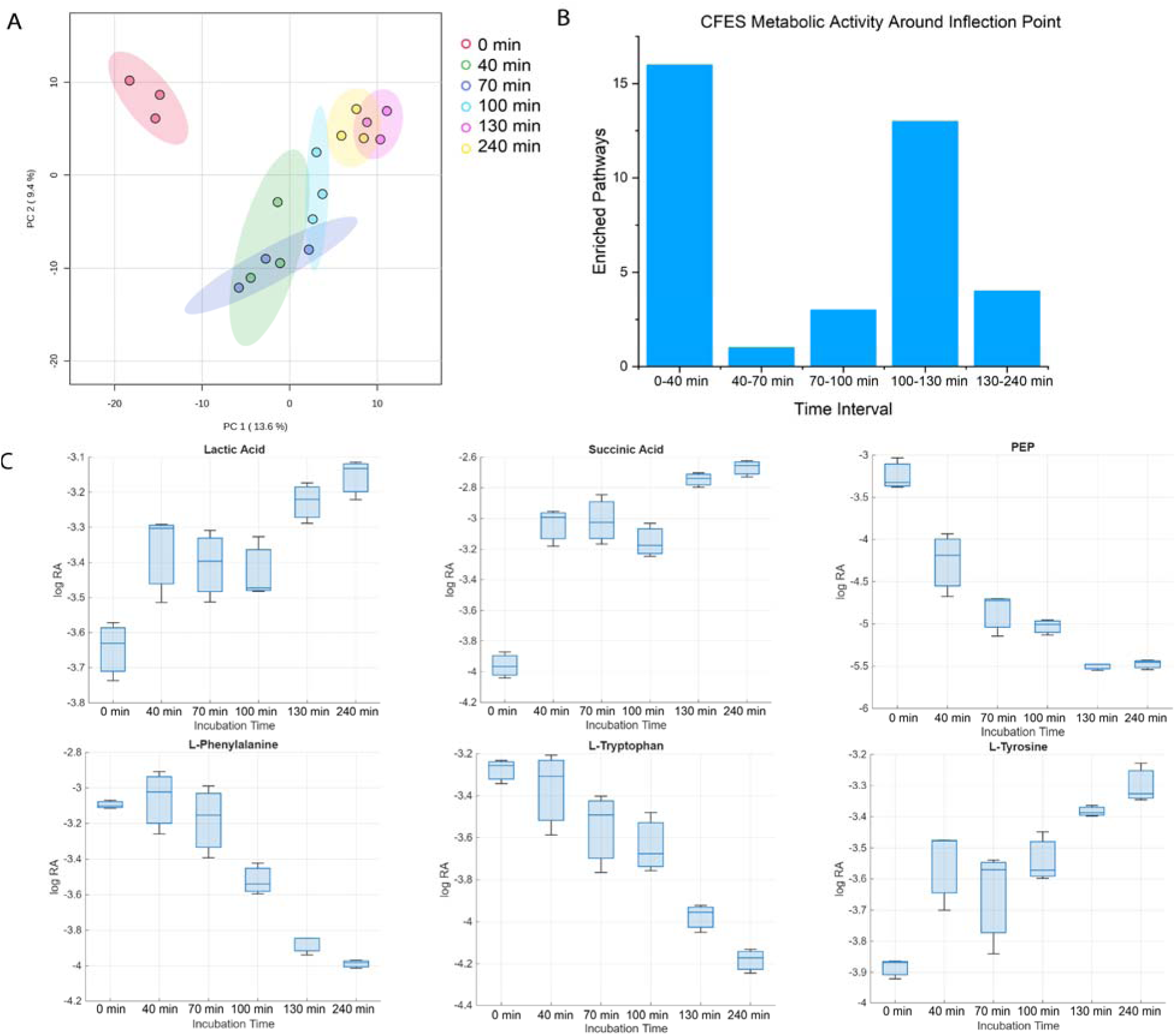
Metabolic characterization around the inflection point of cell-free protein production. sfGFP expression for this run is shown in Figure S1. A) Multivariate analysis of metabolomics results via PCA. Ellipses represent 95% confidence bounds. B) Number of pathways enriched for changes in metabolite levels between consecutive timepoints. Statistical significance for enrichment was determined using a 5% significance level and false discovery rate-corrected p-values. C) Relative abundances of selected individual metabolites, normalized and logarithmically scaled. Blue bars represent medians, boxes represent 25^th^ and 75^th^ percentiles, whiskers indicate extreme values.

Analysis of individual analyte levels can be used to identify some potential biochemical phenomena connected to the changes that CFES undergo as expression rates decrease (Figure 2C). The vast majority of relevant changes happened in metabolites related to energy generation/glycolysis, the TCA cycle, and amino acid metabolism (whether catabolism, biosynthesis, or interconversion). The acceleration of depletion of numerous amino acids (for example, phenylalanine and tryptophan; others are shown in Figure S2) at the inflection point could decrease expression rates not only because amino acids are substrates for translation, but also because they are reactants in metabolic pathways that may impact expression. Based on these results, we sought to identify whether supplementation of cell-free reactions with some of these intermediates might have an impact on either expression or metabolic profiles.

### Analysis of metabolite supplementation and timing

Changes in the levels of numerous amino acids at the inflection point motivated us to test the impact of their supplementation on CFES. While the initial concentration of amino acids in a reaction has been optimized many times over for CFES, these efforts were done before metabolic characterizations revealed the importance and extent of cell-free endogenous metabolism in CFES.^13^ Additionally, these optimizations have typically been done with the assumption that the reaction would be a batch reaction. One exception is that fed-batch supplementation of select amino acids improved expression in the Cytomim expression system, although the timing of the supplementation was arbitrary.^14^ Given that our results suggested that the inflection point is a critical time in cell-free metabolic dynamics, we sought to characterize how supplementing additional amounts of all amino acids (equivalent to their initial concentrations) at different times impacted cell-free expression. For these experiments, we used 10 μL reactions (which have inflection points at 55 min) to simplify experimental logistics.

The impacts of providing reactions with additional amino acids depended heavily on the timing of the supplementation (Figures 3A and 3B). Supplementation after the inflection point led to significant improvements in reaction productivity and lifetime, while supplementation before the inflection point led to significant decreases in both metrics. Moreover, it is the timing of supplementation relative to the inflection point, and not the absolute time, that matters. Adding amino acids at the same absolute time of 75 minutes in two different reaction volumes led to different results (Figure S3): an improvement in expression levels for the 10 μL reaction that has an inflection point of 55 min, and a decrease in performance for the 210 μL reaction that has an inflection point of 100 min. Notably, we were also able to demonstrate that supplementing amino acids after the inflection point also aided protein production at the 210 μL scale. (Figure S4)

**Figure 3:**
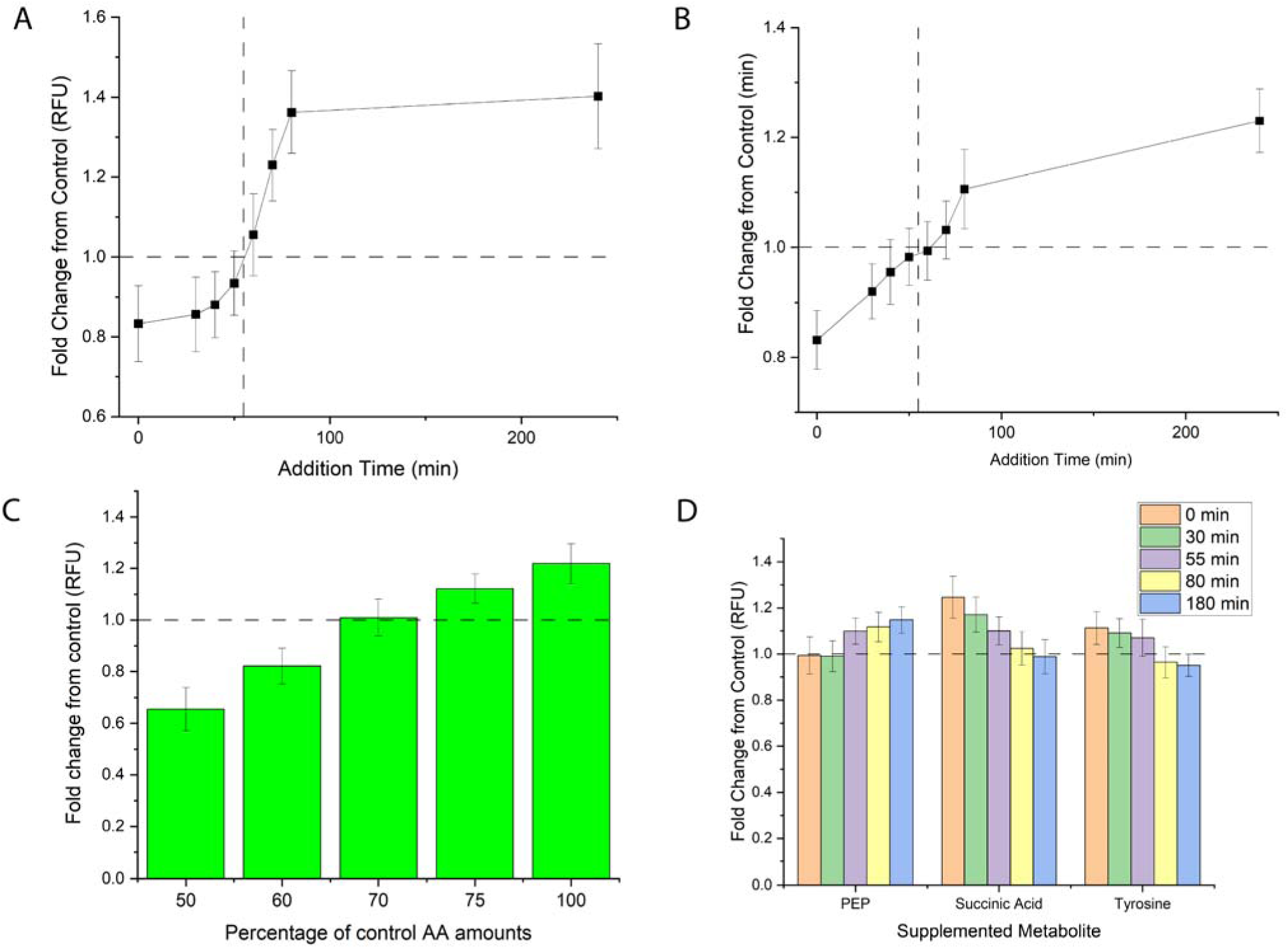
Effect of amino acid supplementation on protein expression in 10 μL CFES reactions. Shown are fold-change improvement (fold change of 1 indicated by horizontal lines) in A) total protein synthesis levels and B) protein synthesis lifetime versus no-supplementation control as a function of the time at which additional amino acids were supplemented, with 55 min being the inflection point time at this reaction volume (indicated by vertical lines). Reactions were supplemented with all amino acids at concentrations equal to their initial concentrations. C) Fold-change improvement in protein synthesis versus a standard reaction control when different levels of total amino acids were added in two steps: half at the beginning of the reaction and half after 120 min. D) Fold-change improvement in total protein synthesis levels versus no-supplementation control as a function of the time at which additional molecules were supplemented at a level equal to their initial concentrations. In all cases, control reactions were supplemented with an equivalent volume of water. All error bars represent ± one standard deviation.

Given that late-reaction supplementation of additional amino acids can improve CFES performance, we next sought to test whether staggering just the initial amino acid levels could provide similar benefits (Figure 3C). We found that splitting the addition of standard amino acid levels between 0 min and 120 min of reaction significantly increased CFES performance (∼21%) compared to addition of those amino acids all at the beginning of the reaction. This finding also led us to test whether similar expression levels could be achieved with less amino acid usage if their addition is staggered across the reaction. We found that amino acid usage could be cut by 30% and still maintain baseline levels of expression if the amino acid addition was staggered in a fed-batch fashion. Taken together, these experiments show that the timing of reagent availability in cell-free reactions can actually be more important to protein production than the total amount of reagent added.

The impact of staggered amino acid addition appears to be due to endogenous metabolic activity rather than simply consumption of amino acids by protein translation. Performing the same experiment in the PURE (Protein synthesis Using Recombinant Elements) expression system, which has minimal endogenous metabolism due to its defined set of components, yielded no significant effect (Figure S5) despite supplementing well after the PURE system’s inflection point in 10 μL reactions of approximately 50 min.^15^ Furthermore, we found that supplementing any individual amino acid well after the inflection point either hurt or did not impact reaction performance (Figure S6). Given that this last observation is different than what was previously reported for the Cytomim system, it further supports the idea that the impacts of supplementation are due to more complex factors than just the exhaustion of specific amino acids by translation.^14^

Fed-batch supplementation of additional quantities of metabolites other than amino acids also had time-dependent impacts on protein expression (Figure 3D). Similar to amino acid supplementation, PEP supplementation after the inflection point improved expression. However, early supplementation did not have negative effects on expression, which is different than what was observed for amino acids and potentially surprising in itself given that inorganic phosphate buildup from energy source usage is known to be an issue in CFES.^16^ In contrast, succinic acid supplementation and tyrosine supplementation had the opposite trend, where early supplementation improved expression but later supplementation did not. Of potential relevance is that both of these metabolites accumulate over the course of a standard cell-free reaction, which is different from the profile seen for most amino acids and thus could be connected to the different impacts of their supplementation timing.

Metabolic characterization of succinic acid supplementation at 0 min revealed different impacts even within the same pathway (Figure S7). Succinic acid profiles remained generally similar, with levels increasing with or without supplementation, though the relative extent of accumulation was less in the supplemented condition. On the other hand, succinic acid supplementation prevented accumulation of several TCA cycle intermediates and instead led to their depletion. Similarly, some non-TCA cycle metabolites also underwent accelerated depletion in the supplemented condition (although the biggest metabolic changes were within the TCA cycle). These observations support the notion that directly perturbing the metabolome in a cell-free reaction can have complex effects on multiple metabolic pathways.

### Metabolic characterization of amino acid supplementation

Metabolic characterization of supplementation of amino acids at 180 min revealed some stark changes in metabolism (Figure 4, additional results in Figure S8). While the absolute differences in PCA scores between the supplemented and control conditions are unsurprising because of the substantial number of metabolites added to the supplemented condition, the metabolic dynamics suggested by PCA are noteworthy. On a systems scale, late-reaction metabolic changes in the supplemented condition appear to be much smaller in magnitude than those seen for reactions without additional amino acids. In addition, the metabolome of the supplemented reaction at 6 hours appears to be much more similar to that of the starting metabolome than is seen with the control condition, at least in the first two principal components. Moreover, supplementation seems to cause several amino acids that normally undergo consumption late in a cell-free reaction to instead maintain roughly constant levels. The exact biochemical mechanisms for this and other supplementation impacts remain unclear.

**Figure 4:**
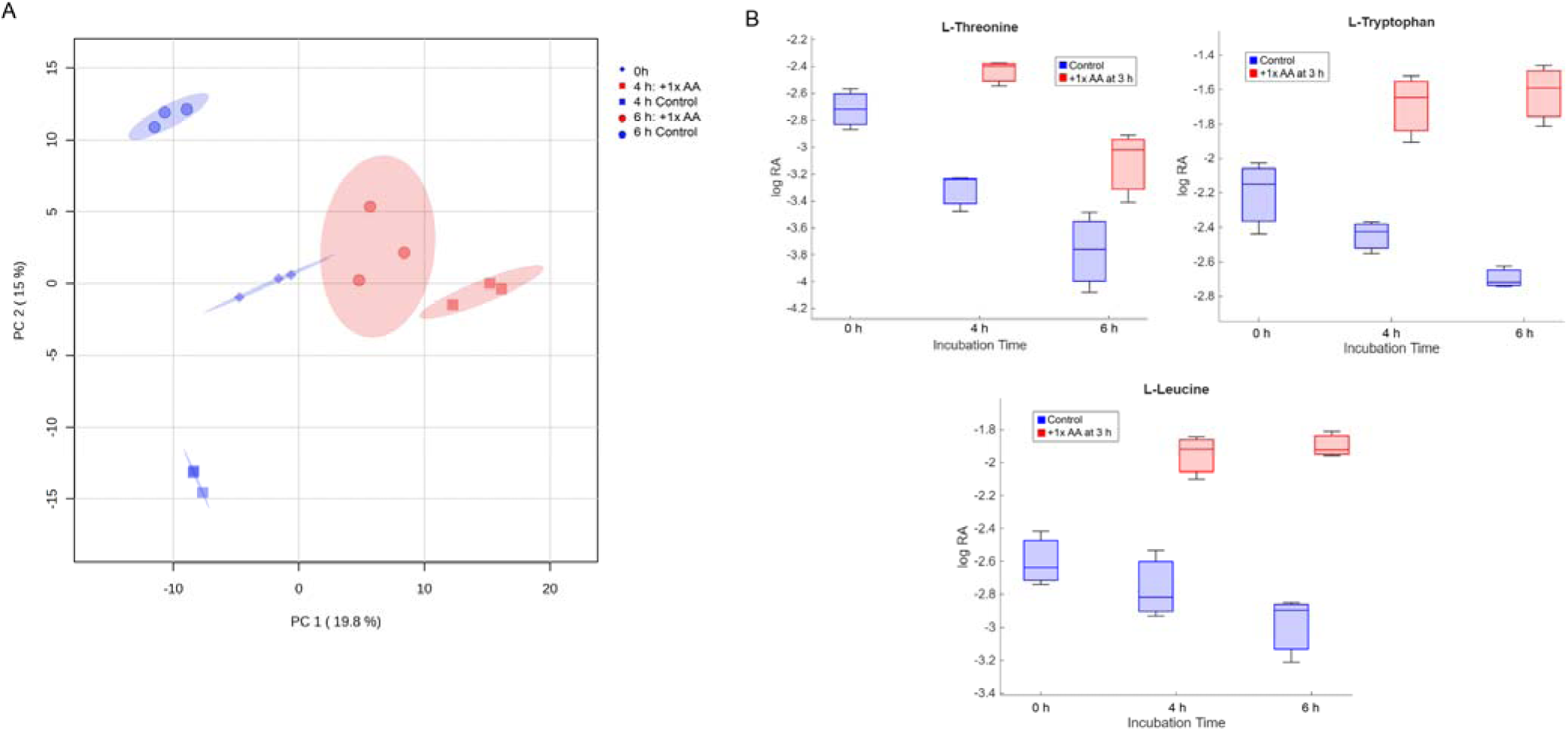
Metabolic impacts of amino acid supplementation at 180 min. A) Multivariate analysis of the metabolome via PCA. Ellipses represent 95% confidence bounds. B) Some individual metabolites with different profiles between the supplemented and non-supplemented conditions. Blue bars represent medians, boxes represent 25^th^ and 75^th^ percentiles, whiskers indicate extreme values. Additional metabolites shown in Figure S8.

In contrast to the apparent decrease in metabolic changes after late supplementation compared to the unsupplemented condition, early supplementation of amino acids at 0 min seems to exaggerate many systems-scale metabolic changes over the course of the reaction (Figure S9). In addition, multiple metabolites have trends in their metabolic profiles that could potentially be linked to negative impacts on expression, from quicker depletion of some amino acids to increased buildup of upper glycolysis metabolites, potentially indicative of less efficient use of energy sources.

## Discussion

In this work, we sought to identify potential connections between two phenomena in CFES. First, some fundamental change in the reaction happens well before protein production stops, which is perhaps most evident in the existence of an inflection point from accelerating to decelerating protein production (Figure 1). Second, metabolism and the metabolic composition of cell-free reactions are known to vary across the course of a reaction and influence protein production. We hypothesized that there may be metabolic changes at the inflection point that could be somehow linked to the end of protein production before stoichiometric theoretical maximum levels.

Metabolomic characterization before and after the inflection point (Figures 2A and 2B) showed a clear systems-scale metabolic shift corresponding with expression slowing down, which in turn suggests the potential importance of metabolism in that deceleration of expression. Like other systems biology approaches, metabolomics analyses are generally limited to identifying correlations rather than inferring causation. Nevertheless, these results can generate testable hypotheses about the mechanisms underlying CFES failure modes. Analyte-level trends suggested that amino acid and energy depletion may contribute to the deceleration of CFES expression, though in likely complex ways. This hypothesis then drove the development of strategies to increase protein production by supplementing the reaction with additional inputs.

These supplementation experiments revealed that for several critical metabolites, not only did the concentration of CFES reaction inputs affect protein expression, but the time at which those inputs were made available to the reaction had a critical impact on reaction performance.

Furthermore, the impact of supplementation timing was clearly connected to the inflection point. For example, adding amino acids before the inflection point hurt both productivity and lifetime, while adding them after the inflection point significantly improved performance (Figure 3).

Moreover, we showed that timing relative to the inflection point was more important than the absolute timescale on which supplementation was performed (Figure S3), and our results suggest that endogenous metabolism was central to these impacts. With previous CFES optimization efforts either using batch supplementation or arbitrarily-timed fed-batch supplementation, the finding that supplementation timing can have a critical impact on protein expression levels opens promising new avenues for cell-free optimization and prototyping.

While the exact biochemical mechanisms for timing-related supplementation impacts remain unclear, metabolic characterization suggested some leads. Fed-batch amino acid supplementation seemed to slow down numerous metabolic changes in the reaction, while doubling initial amino acid levels seemed to exaggerate some metabolic changes. Moreover, the metabolic impacts of supplementation of any given metabolite generally had not just local but also global impacts on metabolism; for example, amino acid supplementation impacted non-amino acid metabolites, and succinic acid supplementation had impacts beyond the TCA cycle. These observations suggest a high degree of interconnectedness in cell-free metabolism even without transcriptional regulation, which in turn suggests the importance of carefully designing and analyzing attempted metabolic interventions or optimization in CFES.

Given that most cell-free reaction additive optimization has been done with batch implementation in mind and with a sparse understanding of cell-free metabolic dynamics, our findings suggest that it is critical to consider the impacts of endogenous metabolism when optimizing the design of reaction components for CFES. Applying our insights to date can help push the yield and scalability of CFES further towards wide industrial application. Furthermore, the importance of endogenous metabolism motivates continuing similar lines of investigation towards a deeper mechanistic understanding of CFES metabolism, which could allow for even greater improvements in CFES performance. In particular, the use of isotope labeling to trace metabolic fluxes could provide more mechanistic insight into specific pathway-level behaviors and limitations driving our observed trends. Resolving endogenous metabolism bottlenecks in CFES would in turn enable more powerful targeted optimization of cell-free systems for synthesis of specific bioproducts, bringing the workflow closer to widespread industrial applicability.

Taken together, our metabolic characterization showed that metabolite supplementation fundamentally alters metabolic dynamics in multiple, complex ways that are dependent on the time of the supplementation. Moreover, these metabolic differences correspond with significant changes in protein expression. These findings provide a foundation for using temporally optimized metabolic interventions to improve the efficiency and productivity of CFES reactions.

## Methods

### Plasmids

The plasmid pJL1 used for the majority of experiments in this study encodes a sfGFP reporter under control of a T7 promoter and strong ribosomal binding site. For studies on inflection time sensitivity pS70s-sfGFP and pS70w-sfGFP were used, which encode sfGFP under control of strong and weak σ70 promoters, respectively.

### Cell-free lysate preparation

Crude extracts consisting of cellular lysates were prepared as described by Sun *et al*.^17^ Briefly, BL21 Star (DE3) *E. coli* cells were grown in 2x YTP media, consisting of 16 g L^−1^ tryptone, 10 g L^−1^ yeast extract, 5 g L^−1^ sodium chloride, 7 g L^−1^ potassium phosphate dibasic, and 3 g L^−1^ potassium phosphate monobasic. Tryptone and yeast extract were purchased from VWR, the rest from Millipore Sigma. Cells were grown at 37°C and 220 rpm in a shaking incubator to the mid-exponential growth phase at an optical density (OD) of 1.5-2.0, with IPTG addition at an OD of 0.6 to induce expression of T7 RNA polymerase. Cells were then centrifuged at 2700 rcf and washed via resuspension with S30A buffer (14 mM magnesium acetate, 60 mM potassium acetate, 10 mM Tris-acetate (pH 8.2), and 2 mM dithiothreitol (DTT), pH-corrected to 7.7 with acetic acid). These centrifugation and resuspension steps were repeated twice, followed by another centrifugation. The wet cell mass was then determined and cells were resuspended in 1 mL of S30A buffer per 1 g of wet cell mass, divided into 1 mL aliquots. Cells were lysed using a Q125 Sonicator (Qsonica, Newton, CT) at a frequency of 20 kHz, and at 50% amplitude. Cells were sonicated in an ice bath with recurring cycles of 10 seconds on and 10 seconds off until approximately 300 J were delivered. An additional 3 mM of DTT was added to each tube, and the sonicated mixture was then centrifuged at 12,000 rcf and 4°C for 10 minutes. The supernatant was carried forward to a runoff reaction. For the runoff reaction, 1 mL aliquots of lysate were incubated in 14 mL round-bottom culture tubes at 37°C and 220 rpm for 80 minutes. The cellular lysate was then centrifuged at 12,000 rcf and 4°C for 10 minutes. The supernatant was removed and loaded into a 10 kDa MWCO dialysis cassette (Thermo Fisher). Lysate was dialyzed in 1L of S30B buffer (14 mM magnesium glutamate, 60 mM potassium glutamate, 1 mM DTT, pH-corrected to 8.2 with Tris base) at 4°C for 3 hours. Dialyzed lysate was removed and centrifuged at 12,000 rcf and 4°C for 10 minutes. The supernatant was removed, aliquoted, and stored at -80°C for future use.

### Cell-free reactions

Cell-free reactions for all experiments were performed as described by Sun *et al*.^17^ Unless otherwise stated, each cell-free reaction contained 0.85 mM each of GTP, UTP, and CTP, in addition to 1.2 mM ATP, 34 μg mL^-1^ of folinic acid, 170 μg mL^-1^ of a commercial *E. coli* tRNA mixture, 130 mM potassium glutamate, 10 mM ammonium glutamate, 12 mM magnesium glutamate, 2 mM of each of the 20 standard amino acids, 0.33 mM nicotine adenine dinucleotide (NAD), 0.27 mM coenzyme-A (CoA), 1.5 mM spermidine, 1 mM putrescine, 4 mM sodium oxalate, 33 mM phosphoenolpyruvate (PEP), 27% cell lysate, and 1 nM of the specified plasmid (except negative controls, which had no plasmid). For each experiment, a fresh aliquot of lysate was used to minimize variability caused by multiple freeze-thaw cycles, and all experiments were performed within three months of lysate preparation. tRNA was purchased from Roche, while all other chemicals used for cell-free reactions were purchased from Millipore Sigma.

For metabolomics analysis, 210 μL reactions were prepared in a 96-well microplate (Falcon, product #351172), which was covered by a breathable film to facilitate gas exchange. Reactions for each time point were aliquoted separately and were run in triplicate, incubated at 37°C for the specified time. 10 μL of the reaction was then removed and stored at -80°C for subsequent fluorescence analysis, and the remaining 200 μL was stored at -80°C for subsequent metabolomics analysis.

In experiments solely assessing GFP production, 10 μL reactions were prepared in triplicate in a 384-well microplate (Grenier Bio-One, product #784101) covered by a breathable film.

Reactions in PURE systems used PURExpress (New England Biolabs) following the manufacturer’s protocol in reaction volumes of 10 μL in triplicate in a 384-well microplate (Grenier Bio-One, product #784101) covered by a breathable film.

### sfGFP measurement

sfGFP fluorescence was measured with a plate reader (Synergy4, BioTek) using excitation and emission wavelengths of 485 and 510 nm, respectively, and a gain of 50.

### Sample processing

Before sample processing, incubated metabolomics samples were used to create a pooled quality control (QC) sample by removing a portion of each sample and combining these aliquots.

Protein precipitation was performed as previously described.^11,12,18^ After samples were thawed, 100% methanol was added to each sample in a 2:1 volumetric ratio to quench the reaction and stop all metabolic activity, followed by incubation at -20°C for at least 15 minutes. Precipitated protein was collected via centrifugation at 11,600 rcf and room temperature for 30 minutes. The resulting supernatant was aliquoted to a microcentrifuge tube and dried using a CentriVap centrifugal concentrator (Labconco) at 40°C until all moisture was removed.

### Gas chromatography-mass spectrometry (GC-MS) analysis

Trimethylsilyl (TMS) derivatization was used to prepare samples for GC-MS analysis as previously described.^11,12,18^ Briefly, 10 μL of a 40 mg/mL solution of methoxyamine hydrochloride (Sigma) in pyridine (Sigma) was added to each sample, which were shaken at 30°C and 1400 rpm for 90 min. Then, 90 μL of N-methyl-N-(trimethylsilyl)-trifluoroacetamide (MSTFA) + 1% trimethylchlorosilane (TMCS) (Thermo Scientific) was added to samples, which were shaken at 37°C and 1400 rpm for 30 min. Samples were centrifuged at 21,100 rcf for 3 min, and 50 μL of the supernatant was added to an autosampler vial with a low-volume insert.

Samples were divided into batches of 12-16, with a QC sample and derivatization blank included in each batch. QCs and derivatization blanks were each injected 3 times per batch. Batches were timed such that the start of a batch matched the end of the previous one, leading to a continuous sequence of injections.

Samples were analyzed by an Agilent 5977/7890b GC-MS with a splitless injection method. The oven temperature was ramped from 50°C to 250°C at a rate of 10°C/min and then held at 250°C for 8 minutes. Additional details on the GC and MS methods are provided in the supplementary information. The column was a Restek Rtx-200 column with an internal diameter of 0.18 mm and a 0.2 μm film thickness.

### Processing and analysis of GC-MS data

Raw GC-MS data files were processed via AMDIS for peak deconvolution.^19^ Peak alignment of conserved peaks was then performed by SpectConnect.^21^ Conserved peaks were putatively annotated by SpectConnect using a previously reported GC-MS library of TMS-derivatized metabolites and a 70% match threshold.^20^

The resulting integrated peak signals underwent preprocessing and imputation via an in-house pipeline. When multiple peaks were annotated as the same metabolite, their integrated signals were summed into a single representative value for that metabolite. Then, metabolites with over 70% missing values (MVs) were eliminated, except that metabolites only present at the start of a cell-free reaction (0 min samples) were retained since we expected there would be a number of analytes that were present at the beginning of the reaction and then quickly depleted. For any given metabolite, conditions (sets of triplicates) with 2 or more MVs had their MVs imputed as small values approximately equal to the smallest measurements in the dataset. The remaining MVs were imputed by no-skip k-nearest neighbors (NS-KNN), an imputation strategy designed for metabolomics data.^22^

MetaboAnalyst 6.0 was used to carry out principal component analysis (PCA), pathway enrichment, and other multivariate analyses, while MATLAB R2025a (MathWorks) was used to plot relative abundance.^23^ A pathway was considered enriched between two conditions if there were multiple metabolite hits and 5% statistical significance using false discovery rate-corrected p-values. Additional details on the rationale for a number of pre-processing choices described above are provided in the supplementary information.

Visualizations were created using Origin Pro (OriginLab)^24^ and BioRender^25^.

## List of abbreviations

CFES: Cell-free expression systems
GC-MS: Gas chromatography-mass spectrometry
MV: Missing value
NS-KNN: No-skip k-nearest neighbor
PCA: Principal component analysis
PEP: Phosphoenolpyruvate
PURE: Protein synthesis using recombinant elements
sfGFP: Superfolder green fluorescent protein
TCA: Tricarboxylic acid

## Declarations

### Ethics approval and consent to participate

Not applicable

### Consent for publication

Not applicable

### Availability of data and materials

The metabolomics datasets used and/or analyzed during this study are available in Metabolomics Workbench (Study ID ST005065).^26^ Other raw data are available from the corresponding author upon request.

### Author Contributions

Conceptualization: S.V. and M.P.S. Investigation: S.V. Formal Analysis: S.V. Writing─Original Draft: S.V. Writing─Review and Editing: S.V. and M.P.S. Visualization: S.V. Supervision: M.P.S. Funding Acquisition: M.P.S.

## Acknowledgements

The plasmid pJL1 was a generous gift from Julius Lucks. The plasmids pS70s-sfGFP and pS70w-sfGFP were a generous gift from Alexandra Patterson.

## References

1. Khanal, O. & Lenhoff, A. M. (2021) Developments and opportunities in continuous biopharmaceutical manufacturing. mAbs vol. 13(1)

2. Hodgman, C. E., & Jewett, M. C. (2012). Cell-free synthetic biology: Thinking outside the cell. Metabolic Engineering, 14(3), 261–269.

3. Zhang, Y., Steppe, P. L., Kazman, M. W., & Styczynski, M. P. (2021). Point-of-Care Analyte Quantification and Digital Readout via Lysate-Based Cell-Free Biosensors Interfaced with Personal Glucose Monitors. ACS Synthetic Biology, 10(11), 2862–2869.

4. McSweeney, M. A., Patterson, A. T., Loeffler, K., Cuellar Lelo De Larrea, R., McNerney, M. P., Kane, R. S., & Styczynski, M. P. (2025). A modular cell-free protein biosensor platform using split T7 RNA polymerase. Sci. Adv (Vol. 11).

5. Sitaraman, K., Esposito, D., Klarmann, G., le Grice, S. F., Hartley, J. L., & Chatterjee, D. K. (2004). A novel cell-free protein synthesis system. Journal of Biotechnology, 110(3), 257–263.

6. Batista AC, Soudier P, Kushwaha M, Faulon JL. (2021) Optimising protein synthesis in cell free systems, a review. *Eng*. Biol., 5:10–19.

7. Dopp, J. L., & Reuel, N. F. (2018). Process optimization for scalable E. coli extract preparation for cell-free protein synthesis. Biochemical Engineering Journal, 138, 21–28.

8. Caschera, F., & Noireaux, V. (2014). Synthesis of 2.3 mg/ml of protein with an all Escherichia coli cell-free transcription-translation system. Biochimie, 99(1), 162–168.

9. Johnson, C., Ivanisevic, J. & Siuzdak, G. Metabolomics: beyond biomarkers and towards mechanisms. Nat Rev Mol Cell Biol 17, 451–459 (2016).

10. Somvanshi, P. R., & Venkatesh, K. V. (2014). A conceptual review on systems biology in health and diseases: From biological networks to modern therapeutics. Systems and Synthetic Biology, 8(1), 99–116.

11. Miguez, A. M., McNerney, M. P., & Styczynski, M. P. (2019). Metabolic Profiling of Escherichia coli-Based Cell-Free Expression Systems for Process Optimization. Industrial and Engineering Chemistry Research, 58(50), 22472–22482. 10.1021/acs.iecr.9b03565

12. Miguez, A. M., Zhang, Y., Piorino, F., & Styczynski, M. P. (2021). Metabolic Dynamics in Escherichia coli-Based Cell-Free Systems. ACS Synthetic Biology, 10(9), 2252–2265.

13. Kim, D. M., & Swartz, J. R. (2001). Regeneration of adenosine triphosphate from glycolytic intermediates for cell-free protein synthesis. Biotechnology and Bioengineering, 74(4), 309–316.

14. Jewett, M. C., Calhoun, K. A., Voloshin, A., Wuu, J. J., & Swartz, J. R. (2008). An integrated cell-free metabolic platform for protein production and synthetic biology. Molecular Systems Biology, 4.

15. Shimizu, Y., Kuruma, Y., Kanamori, T., & Ueda, T. (2014). The PURE system for protein production. Methods in Molecular Biology, 1118, 275–284.

16. Calhoun KA, Swartz JR. Energy systems for ATP regeneration in cell-free protein synthesis reactions. (2014) Methods in Molecular Biology. 375:3–17.

17. Sun, Z. Z., Hayes, C. A., Shin, J., Caschera, F., Murray, R. M., & Noireaux, V. (2013). Protocols for implementing an Escherichia coli based TX-TL cell-free expression system for synthetic biology. Journal of Visualized Experiments, 79.

18. Miguez, A. M., Zhang, Y., & Styczynski, M. P. (2022). Metabolomics Analysis of Cell-Free Expression Systems Using Gas Chromatography-Mass Spectrometry. In Methods in Molecular Biology (Vol. 2433, pp. 217–226). Humana Press Inc.

19. Mallard, W. G., & Stein, S. E. (1997). An integrated method for spectrum extraction and compound identification from GC/MS data. Journal of the American Society for Mass Spectrometry, 8(3), 239–248.

20. Kind, T., Wohlgemuth, G., Yup Lee, D., Palazoglu, M., Shahbaz, S., Fiehn, O. (2009). FiehnLib: Mass Spectral and Retention Index Libraries for Metabolomics Based on Quadrupole and Time-of-Flight Gas Chromatography/Mass Spectrometry. Analytical Chemistry, 2009.

21. Styczynski, M. P., Moxley, J. F., Tong, L. v., Walther, J. L., Jensen, K. L., & Stephanopoulos, G. N. (2007). Systematic identification of conserved metabolites in GC/MS data for metabolomics and biomarker discovery. Analytical Chemistry, 79(3), 966–973.

22. Lee, J. Y., & Styczynski, M. P. (2018). NS-kNN: a modified k-nearest neighbors approach for imputing metabolomics data. Metabolomics, 14(12). 10.1007/s11306-018-1451-8

23. Pang, Z., Lu, Y., Zhou, G., Hui, F., Xu, L., Viau, C., Spigelman, A. F., Macdonald, P. E., Wishart, D. S., Li, S., & Xia, J. (2024). MetaboAnalyst 6.0: towards a unified platform for metabolomics data processing, analysis and interpretation. Nucleic Acids Research, 52(W1), W398–W406.

24. Origin(Pro), Version 2025, OriginLab Corporation, Northampton, MA, USA.

25. BioRender.com. (2025). Retrieved from https://biorender.com

26. Sud M, Fahy E, Cotter D, Azam K, Vadivelu I, Burant C, Edison A, Fiehn O, Higashi R, Nair KS, Sumner S, Subramaniam S. (2016) Metabolomics Workbench: An international repository for metabolomics data and metadata, metabolite standards, protocols, tutorials and training, and analysis tools. Nucleic Acids Research. 44(D1)

