## Supplementary Figures for "Timing of metabolomics-driven supplementation strategies affects protein expression in *E. coli-*based cell-free expression systems"

**Corresponding Author:**

### GC-MS Methods

*Auto-sampler Method*

| Solvent | Injection Volume (μL) | Samples per derivatization batch | Solvent washes |
|---|---|---|---|
| Pyridine | 1 | 12-18 | 3 pre-injection, 3 post-injection |

*GC Method*

| Initial Oven Temperature (°C) | Final Oven Temperature (°C) | Ramp Rate (°C/min) | Final Temperature Hold (min) | Column Flow (mL/min) | Total Flow (mL/min) |
|---|---|---|---|---|---|
| 50 | 305 | 10 | 8 | 1 | 20 |

*MS Method*

| Ion Source Temperature (°C) | Quadrupole Temperature (°C) | Scanning Range (m/z) | MS Transfer Line Temperature (°C) |
|---|---|---|---|
| 230 | 150 | 50-500 | 310 |

*Rationale for selected data processing choices*

| Methodology | Rationale |
|---|---|
| Constant missing value replacement with 2/3 replicate threshold | Metabolites with 2/3 or 3/3 values missing in a given condition should be considered absent and replaced with an arbitrarily small value. Replacement is done with half of the lowest integrated signal of the given metabolite (plus noise) to best approximate going below LOD. |
| NS-KNN with k of 2 | For metabolites with just 1/3 missing values in a given condition, the metabolite should be treated as present and imputed using the two mathematically closest samples (likely the other two replicates). Additional |

| | |
|---|---|
| | analysis indicated minimal sensitivity of results to increasing k values beyond 2 (data not shown). |
| Data filtering based on missing values (>70%) | Metabolites with more than 70% missing values will be contributing more noise than signal to the dataset, potentially confounding the ability to make multivariate interpretations. |

**Plasmids used in this study:**

All plasmids used in this study contain a high copy number colE1 origin of replication and a kanamycin resistance cassette controlled by a constitutive endogenous promoter. Sequences are shown below, with the sfGFP expression cassette capitalized and highlighted in green:

pJL1 (2486 bp, under strong T7 promoter):

**agatcaaaggatcttcttgagatcctttttttctgcgcgtaatctgctgcttgcaaacaaaaaaaccaccgctaccagcggtggtttgt
ttgccggatcaagagctaccaactctttttccgaaggtaactggcttcagcagagcgcagataccaaatactgttcttctagtgtagc
cgtagttaggccaccacttcaagaactctgtagcaccgcctacatacctcgctctgctaatcctgttaccagtggctgctgccagtgg
cgataagtcgtgtcttaccgggttggactcaagacgatagttaccggataaggcgcagcggtcgggctgaacggggggttcgtgca
cacagcccagcttggagcgaacgacctacaccgaactgagatacctacagcgtgagctatgagaaagcgccacgcttcccgaag
ggagaaaggcggacaggtatccggtaagcggcagggtcggaacaggagagcgcacgagggagcttccagggggaaacgcctg
gtatctttatagtcctgtcgggtttcgccacctctgacttgagcgtcgatttttgtgatgctcgtcaggggggcggagcctatggaaaa
acgccagcaacgcgatcccgcgaaattaatacgactcactatagggagaccacaacggtttccctctagaaataattttgtttaact
ttaagaaggagatatacatATGAGCAAAGGTGAAGAACTGTTTACCGGCGTTGTGCCGATT
CTGGTGGAACTGGATGGCGATGTGAACGGTCACAAATTCAGCGTGCGTGGTGA**

**AGGTGAAGGCGATGCCACGATTGGCAAACTGACGCTGAAATTTATCTGCACCAC
CGGCAAACTGCCGGTGCCGTGGCCGACGCTGGTGACCACCCTGAfCCTATGGC
GTTCAGTGTTTTAGTCGCTATCCGGATCACATGAAACGTCACGATTTCTTTAAAT
CTGCAATGCCGGAAGGCTATGTGCAGGAACGTACGATTAGCTTTAAAGATGATG
GCAAATATAAAACGCGCGCCGTTGTGAAATTTGAAGGCGATACCCTGGTGAACC
GCATTGAACTGAAAGGCACGGATTTTAAAGAAGATGGCAATATCCTGGGCCATA
AACTGGAATACAACTTTAATAGCCATAATGTTTATATTACGGCGGATAAACAGAA
AAATGGCATCAAAGCGAATTTTACCGTTCGCCATAACGTTGAAGATGGCAGTGT
GCAGCTGGCAGATCATTATCAGCAGAATACCCCGATTGGTGATGGTCCGGTGCT
GCTGCCGGATAATCATTATCTGAGCACGCAGACCGTTCTGTCTAAAGATCCGAA
CGAAAAAGGCACGCGGGACCACATGGTTCTGCACGAATATGTGAATGCGGCAG
GTATTACGTGGAGCCATCCGCAGTTCGAAAAATAAgtcgaccggctgctaacaaagcccgaaagg
aagctgagttggctgctgccaccgctgagcaataactagcataaccccttggggcctctaaacgggtcttgaggggttttttgctgaa
agccaattctgattagaaaaactcatcgagcatcaaatgaaactgcaatttattcatatcaggattatcaataccatatttttgaaaa
agccgtttctgtaatgaaggagaaaactcaccgaggcagttccataggatggcaagatcctggtatcggtctgcgattccgactcgt
ccaacatcaatacaacctattaatttcccctcgtcaaaaataaggttatcaagtgagaaatcaccatgagtgacgactgaatccggt
gagaatggcaaaagcttatgcatttctttccagacttgttcaacaggccagccattacgctcgtcatcaaaatcactcgcatcaacca
aaccgttattcattcgtgattgcgcctgagcgagacgaaatacgcgatcgctgttaaaaggacaattacaaacaggaatcgaatgc
aaccggcgcaggaacactgccagcgcatcaacaatattttcacctgaatcaggatattcttctaatacctggaatgctgttttcccgg
ggatcgcagtggtgagtaaccatgcatcatcaggagtacggataaaatgcttgatggtcggaagaggcataaattccgtcagcca
gtttagtctgaccatctcatctgtaacatcattggcaacgctacctttgccatgtttcagaaacaactctggcgcatcggcttcccat
acaatcgatagattgtcgcacctgattgcccgacattatcgcgagcccatttatacccatataaatcagcatccatgttggaatttaat**

**cgcggcttcgagcaagacgtttcccgttgaatatggctcataacaccccttgtattactgtttatgtaagcagacagttttattgttcat**

**gatgatatatttttatcttgtgcaatgtaacatcagagattttgagacacaacgtg**

pS70s-sfGFP (3375 bp, under strong $\sigma^{70}$ promoter):

**gcgctagcggagtgtatactggcttactatgttggcactgatgagggtgtcagtgaagtgcttcatgtggcaggagaaaaaaggct**

**gcaccggtgcgtcagcagaatatgtgatacaggatatattccgcttcctcgctcactgactcgctacgctcggtcgttcgactgcggc**

**gagcggaaatggcttacgaacggggcggagatttcctggaagatgccaggaagatacttaacagggaagtgagagggccgcgg**

**caaagccgtttttccataggctccgcccccctgacaagcatcacgaaatctgacgctcaaatcagtggtggcgaaacccgacagga**

**ctataaagataccaggcgtttccccctggcggctccctcgtgcgctctcctgttcctgcctttcggtttaccggtgtcattccgctgttat**

**ggccgcgtttgtctcattccacgcctgacactcagttccgggtaggcagttcgctccaagctggactgtatgcacgaaccccccgttc**

**agtccgaccgctgcgccttatccggtaactatcgtcttgagtccaacccggaaagacatgcaaaagcaccactggcagcagccact**

**ggtaattgatttagaggagttagtcttgaagtcatgcgccggttaaggctaaactgaaaggacaagttttggtgactgcgctcctcca**

**agccagttacctcggttcaaagagttggtagctcagagaaccttcgaaaaaccgccctgcaaggcggttttttcgttttcagagcaa**

**gagattacgcgcagaccaaaacgatctcaagaagatcatcttattaatcagataaaatatttctagatttcagtgcaatttatctcttc**

**aaatgtagcacctgaagtcagccccatacgatataagttgtaattctcatgtttgacagcttatcatcgataagcttccgatggcgcg**

**ccgagaggctttacactttatgcttccggctgaattctaaagatctttgacagctagctcagtcctaggtataatactagtacgtcgac**

**tctcgagtgagattgttgacggtaccgtattttggatctaggaggaaggatctATGAGCAAAGGAGAAGAACTT**

**TTCACTGGAGTTGTCCCAATTCTTGTTGAATTAGATGGTGATGTTAATGGGCACA**

**AATTTTCTGTCCGTGGAGAGGGTGAAGGTGATGCTACAAACGGAAAACTCACCC**

**TTAAATTTATTTGCACTACTGGAAAACTACCTGTTCCGTGGCCAACACTTGTCAC**

**TACTCTGACCTATGGTGTTCAATGCTTTTCCCGTTATCCGGATCACATGAAACGG**

**CATGACTTTTTCAAGAGTGCCATGCCCGAAGGTTATGTACAGGAACGCACTATAT**

**CTTTCAAAGATGACGGGACCTACAAGACGCGTGCTGAAGTCAAGTTTGAAGGT**

**GATACCCTTGTTAATCGTATCGAGTTAAAGGGTATTGATTTTAAAGAAGATGGAA**

**ACATTCTTGGACACAAACTCGAGTACAACTTTAACTCACACAATGTATACATCAC**

**GGCAGACAAACAAAAGAATGGAATCAAAGCTAACTTCAAAATTCGCCACAACGT**

**TGAAGATGGTTCCGTTCAACTAGCAGACCATTATCAACAAAATACTCCAATTGGC**

**GATGGCCCTGTCCTTTTACCAGACAACCATTACCTGTCGACACAATCTGTCCTTT**

**CGAAAGATCCCAACGAAAAGCGTGACCACATGGTCCTTCTTGAGTTTGTAACTG**

**CTGCTGGGATTACACATGGCATGGATGAGCTCTACAAAtaaggatctgaagcttgggcccgaac**

**aaaaactcatctcagaagaggatctgaatagcgccgtcgaccatcatcatcatcatcattgagtttaaacggtctccagcttggctgt**

**tttggcggatgagagaagattttcagcctgatacagattaaatcagaacgcagaagcggtctgataaaacagaatttgcctggcgg**

**cagtagcgcggtggtcccacctgaccccatgccgaactcagaagtgaaacgccgtagcgccgatggtagtgtggggtctccccatg**

**cgagagtagggaactgccaggcatcaaataaaacgaaaggctcagtcgaaagactgggcctttcgttttatctgttgtttgtcggtg**

**aactggatccttactcgagtctagactgcagttgatcgggcacgtaagaggttccaactttcaccataatgaaataagatcactacc**

**gggcgtatttttttgagttatcgagattttcaggagctaaggaagctaaaatggagaaaaaaatcactggatataccaccgttgatat**

**atcccaatggcatcgtaaagaacattttgaggcatttcagtcagttgctcaatgtacctataaccagaccgttcagctggatattacg**

**gcctttttaaagaccgtaaagaaaaataagcacaagttttatccggcctttattcacattcttgcccgcctgatgaatgctcatccgga**

**atttcgtatggcaatgaaagacggtgagctggtgatatgggatagtgttcacccttgttacaccgttttccatgagcaaactgaaacg**

**ttttcatcgctctggagtgaataccacgacgatttccggcagtttctacacatatattcgcaagatgtggcgtgttacggtgaaaacct**

**ggcctatttccctaaagggtttattgagaatatgtttttcgtctcagccaatccctgggtgagtttcaccagttttgatttaaacgtggcc**

**aatatggacaacttcttcgcccccgttttcaccatgggcaaatattatacgcaaggcgacaaggtgctgatgccgctggcgattcag**

**gttcatcatgccgtttgtgatggcttccatgtcggcagaatgcttaatgaattacaacagtactgcgatgagtggcagggcggggcg**

**taatttgatatcgagctcgcttggactcctgttgatagatccagtaatgacctcagaactccatctggatttgttcagaacgctcggtt**

**gccgccgggcgtttttttattggtgagaatccaagcctccgatcaacgtctcattttcgccaaaagttggcccagggcttcccggtatc**

**aacagggacaccaggatttatttattctgcgaagtgatcttccgtcacaggtatttattcggcgcaaagtgcgtcgggtgatgctgcc**

**aacttactgatttagtgtatgatggtgtttttgaggtgctccagtggcttctgtttctatcagctgtccctcctgttcagctactgacggg gtggtgcgtaacggcaaaagcaccgccggacaaatca**

pS70w-sfGFP (2498 bp, under weak $\sigma^{70}$ promoter):

**agatcaaaggatcttcttgagatcctttttttctgcgcgtaatctgctgcttgcaaacaaaaaaaccaccgctaccagcggtggtttgt ttgccggatcaagagctaccaactctttttccgaaggtaactggcttcagcagagcgcagataccaaatactgttcttctagtgtagc cgtagttaggccaccacttcaagaactctgtagcaccgcctacatacctcgctctgctaatcctgttaccagtggctgctgccagtgg cgataagtcgtgtcttaccgggttggactcaagacgatagttaccggataaggcgcagcggtcgggctgaacggggggttcgtgca cacagcccagcttggagcgaacgacctacaccgaactgagatacctacagcgtgagctatgagaaagcgccacgcttcccgaag ggagaaaggcggacaggtatccggtaagcggcagggtcggaacaggagagcgcacgagggagcttccagggggaaacgcctg gtatctttatagtcctgtcgggtttcgccacctctgacttgagcgtcgatttttgtgatgctcgtcaggggggcggagcctatggaaaa acgccagcaacgcgatcccgcgaaattttacggctagctcagtcctaggtactatgctagcccacaacggtttccctctagaaataa ttttgtttaactttaagaaggagatatacatATGAGCAAAGGTGAAGAACTGTTTACCGGCGTTGT GCCGATTCTGGTGGAACTGGATGGCGATGTGAACGGTCACAAATTCAGCGTGC GTGGTGAAGGTGAAGGCGATGCCACGATTGGCAAACTGACGCTGAAATTTATCT GCACCACCGGCAAACTGCCGGTGCCGTGGCCGACGCTGGTGACCACCCTGACC TATGGCGTTCAGTGTTTTAGTCGCTATCCGGATCACATGAAACGTCACGATTTCT TTAAATCTGCAATGCCGGAAGGCTATGTGCAGGAACGTACGATTAGCTTTAAAG ATGATGGCAAATATAAAACGCGCGCCGTTGTGAAATTTGAAGGCGATACCCTGG TGAACCGCATTGAACTGAAAGGCACGGATTTTAAAGAAGATGGCAATATCCTGG GCCATAAACTGGAATACAACTTTAATAGCCATAATGTTTATATTACGGCGGATAAA CAGAAAAATGGCATCAAAGCGAATTTTACCGTTCGCCATAACGTTGAAGATGGC AGTGTGCAGCTGGCAGATCATTATCAGCAGAATACCCCGATTGGTGATGGTCCG GTGCTGCTGCCGGATAATCATTATCTGAGCACGCAGACCGTTCTGTCTAAAGAT**

**CCGAACGAAAAAGGCACGCGGGACCACATGGTTCTGCACGAATATGTGAATGC**

**GGCAGGTATTACGTGGAGCCATCCGCAGTTCGAAAAATAAgtcgaccggctgctaacaaagc**

**ccgaaaggaagctgagttggctgctgccaccgctgagcaataactagcataaccccttggggcctctaaacgggtcttgaggggtt**

**ttttgctgaaagccaattctgattagaaaaactcatcgagcatcaaatgaaactgcaatttattcatatcaggattatcaataccatat**

**ttttgaaaaagccgtttctgtaatgaaggagaaaactcaccgaggcagttccataggatggcaagatcctggtatcggtctgcgatt**

**ccgactcgtccaacatcaatacaacctattaatttcccctcgtcaaaaataaggttatcaagtgagaaatcaccatgagtgacgact**

**gaatccggtgagaatggcaaaagcttatgcatttctttccagacttgttcaacaggccagccattacgctcgtcatcaaaatcactcg**

**catcaaccaaaccgttattcattcgtgattgcgcctgagcgagacgaaatacgcgatcgctgttaaaaggacaattacaaacagga**

**atcgaatgcaaccggcgcaggaacactgccagcgcatcaacaatattttcacctgaatcaggatattcttctaatacctggaatgct**

**gttttcccggggatcgcagtggtgagtaaccatgcatcatcaggagtacggataaaatgcttgatggtcggaagaggcataaattc**

**cgtcagccagtttagtctgaccatctcatctgtaacatcattggcaacgctacctttgccatgtttcagaaacaactctggcgcatcgg**

**gcttcccatacaatcgatagattgtcgcacctgattgcccgacattatcgcgagcccatttatacccatataaatcagcatccatgttg**

**gaatttaatcgcggcttcgagcaagacgtttcccgttgaatatggctcataacaccccttgtattactgtttatgtaagcagacagtttt**

**attgttcatgatgatatatttttatcttgtgcaatgtaacatcagagattttgagacacaacgtg**

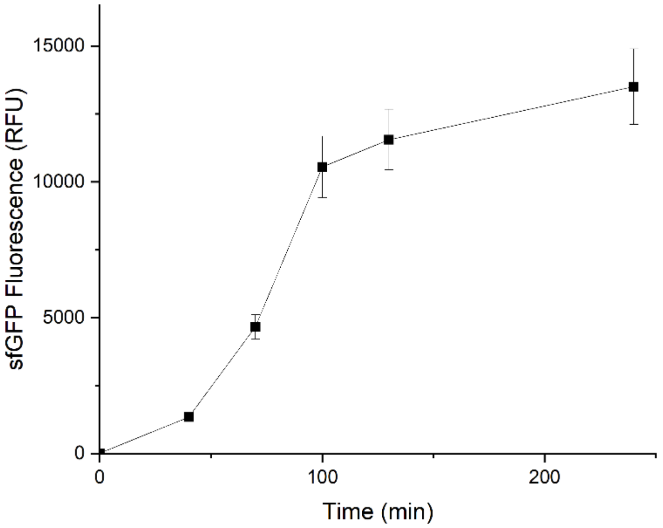


**Figure S1:** Time course expression profile corresponding to the metabolomics results in Figure 2. Error bars indicate ± one standard deviation. As expected, expression slows down after 100 min.

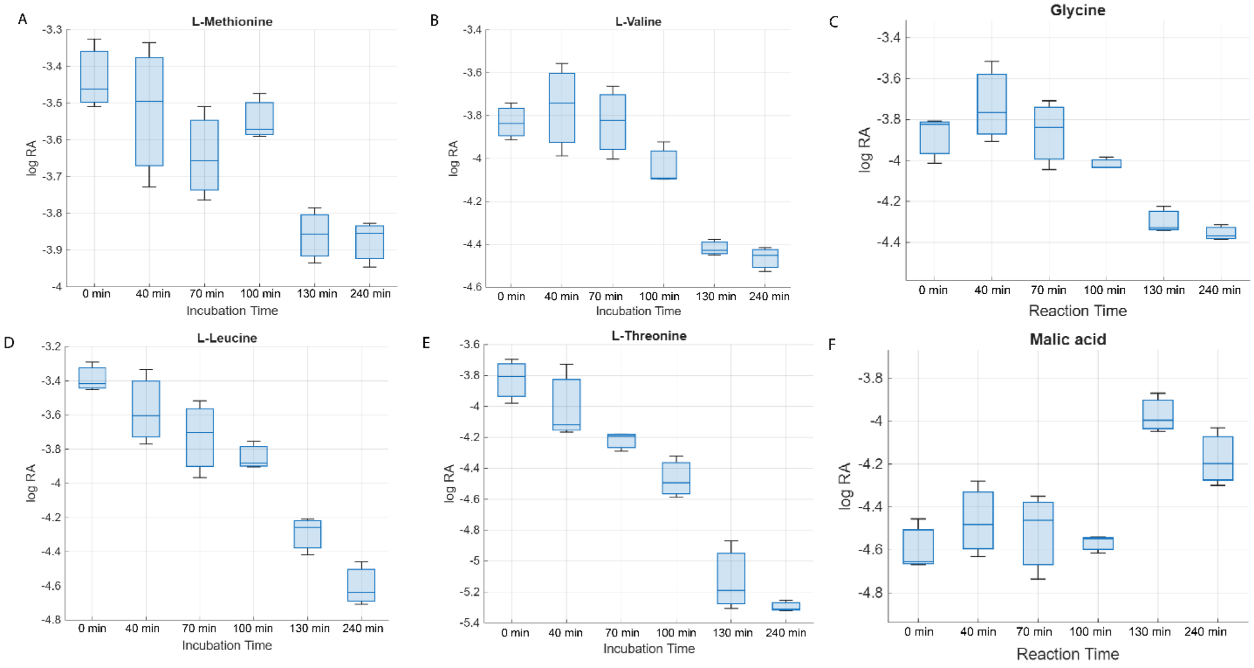


**Figure S2:** Additional analyte profiles for the inflection point study shown in Figure 3: A) L-methionine, B) L-valine, C) glycine, D) L-leucine, E) L-threonine, F) Malic acid. Blue bars represent medians, boxes represent 25$^{th}$ and 75$^{th}$ percentiles, whiskers indicate extreme values.

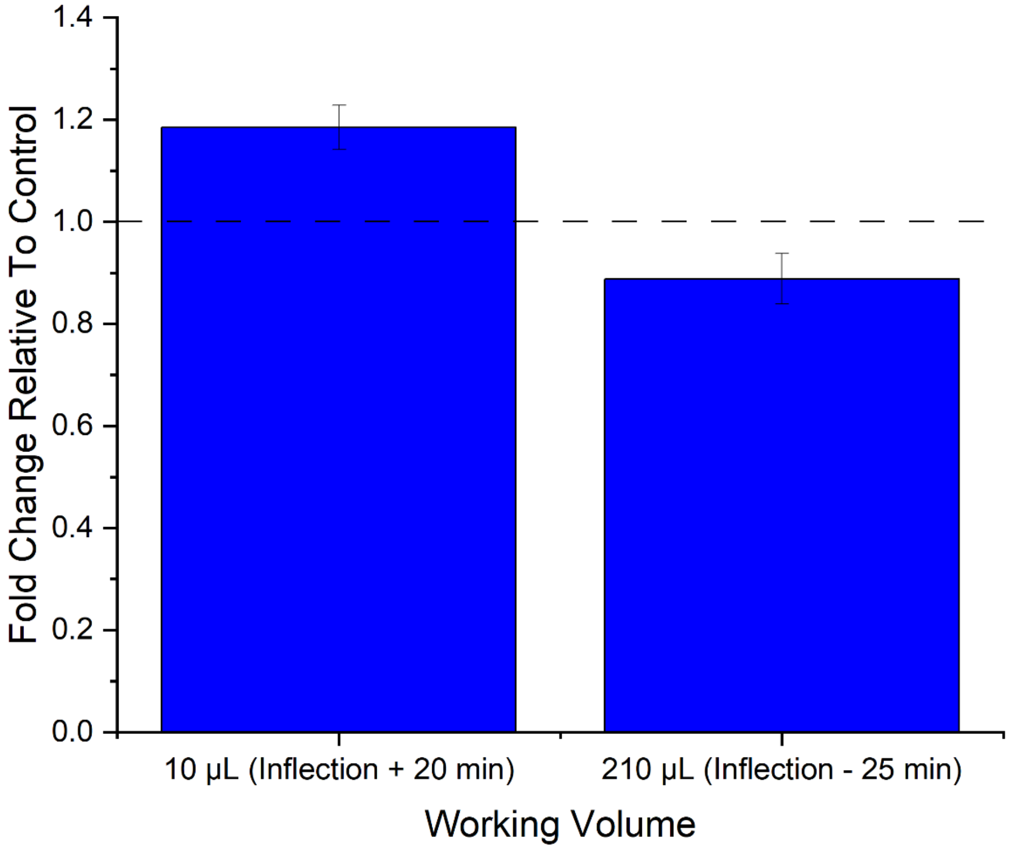


**Figure S3:** Effect of supplementing additional amino acids at 75 min for two different reaction volumes. At a working volume of 10 µL, the supplementation time is 20 minutes after the inflection point and leads to increased expression. At a working volume of 210 µL, the supplementation time is 25 minutes before the inflection point and leads to decreased expression. Error bars represent ± one standard deviation of propagated error.

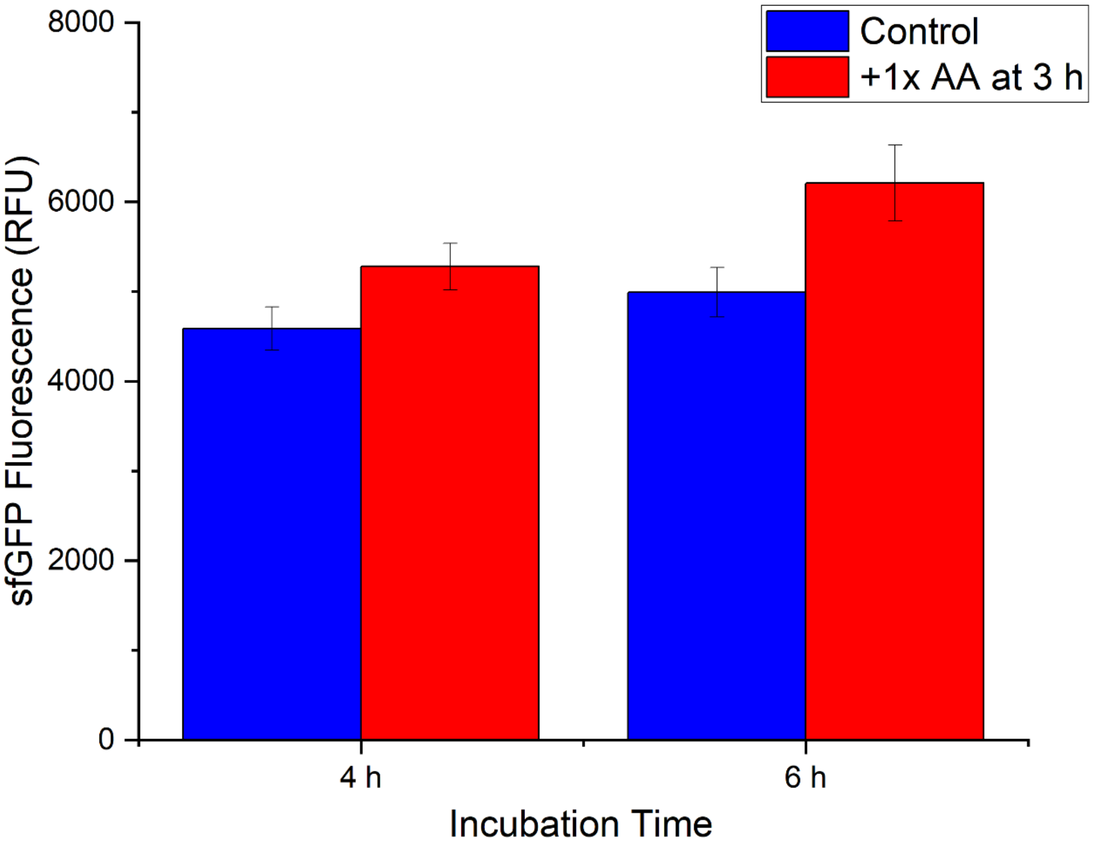


**Figure S4:** Effect of supplementing additional amino acids at 3 h in a reaction volume of 210 μL. Similar to the smaller-scale 10 μL results shown in Figure S3, post-inflection point supplementation improves CFES protein production. Error bars represent ± one standard deviation.

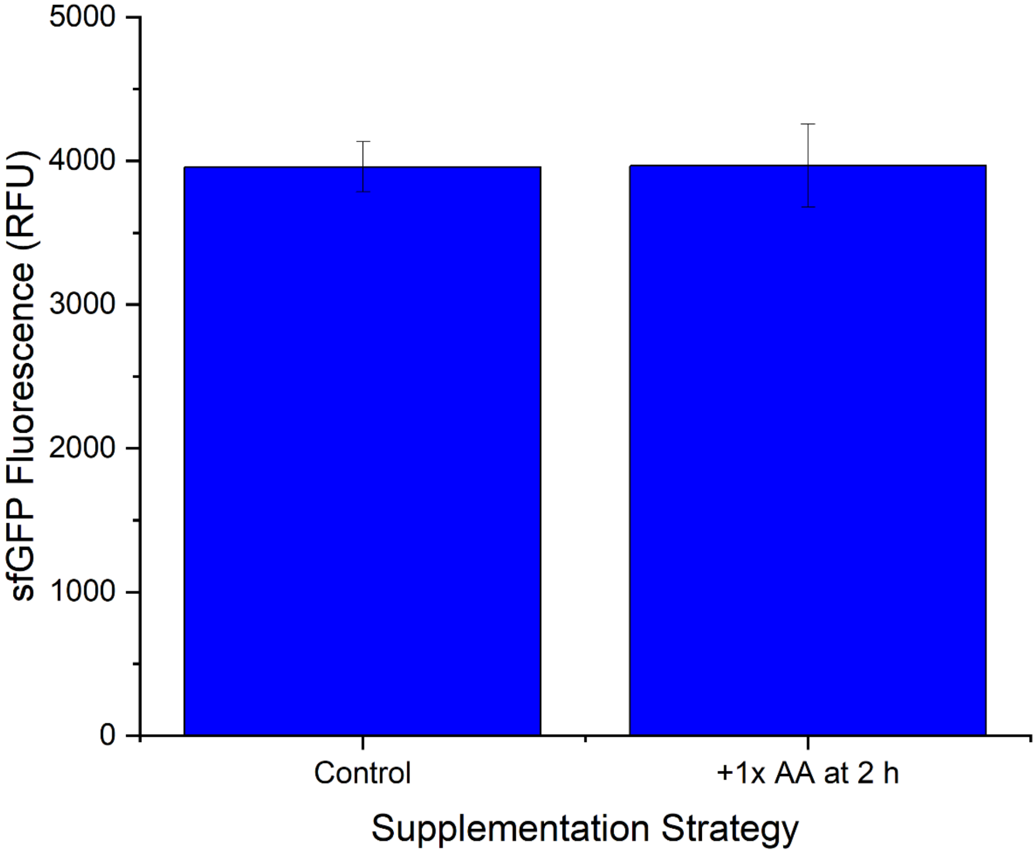


**Figure S5:** Impact of supplementing additional amino acids in a PURE system after 2 h of reaction. The inflection point in this PURE cell-free system in 10 µL reaction volumes occurred at approximately 50 min. Adding amino acids after the inflection point had no significant effect, supporting the idea that the post-inflection point supplementation impacts in lysate based-systems are linked to endogenous metabolism in the lysate. Error bars represent ± one standard deviation.

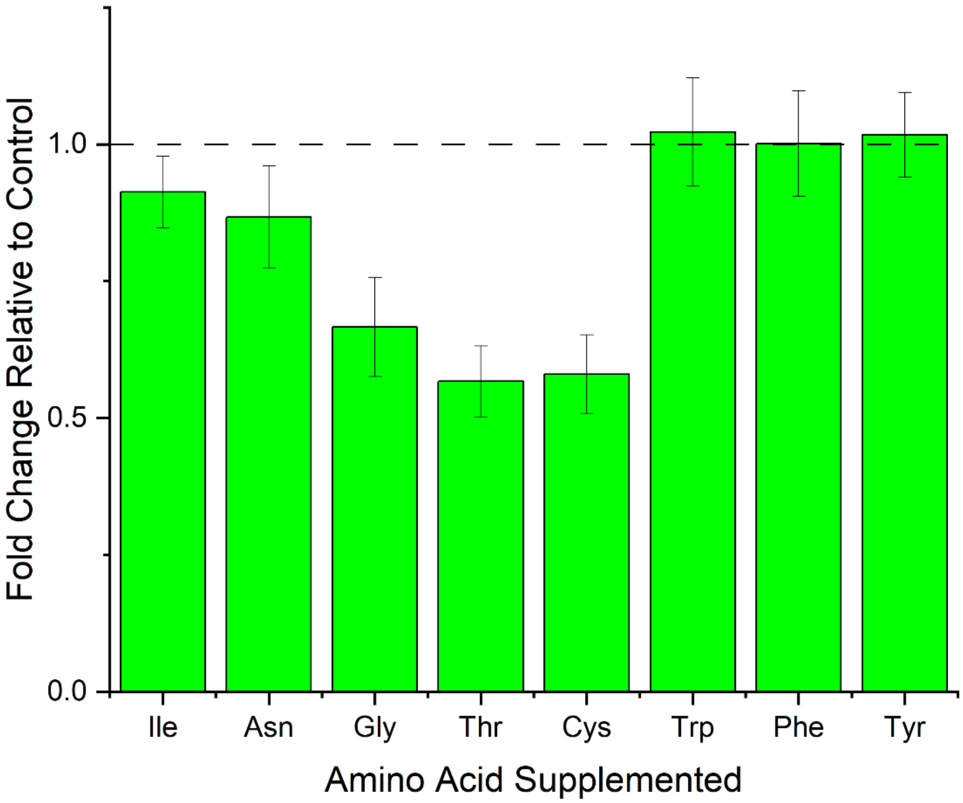


**Figure S6:** Effect of supplementing individual amino acids after the inflection point (at 240 min) on CFES productivity. Unlike supplementation of all amino acids, supplementing 2 mM of any individual amino acid had either no discernable effect or negative impacts on productivity. Error bars represent ± one standard deviation of propagated error.

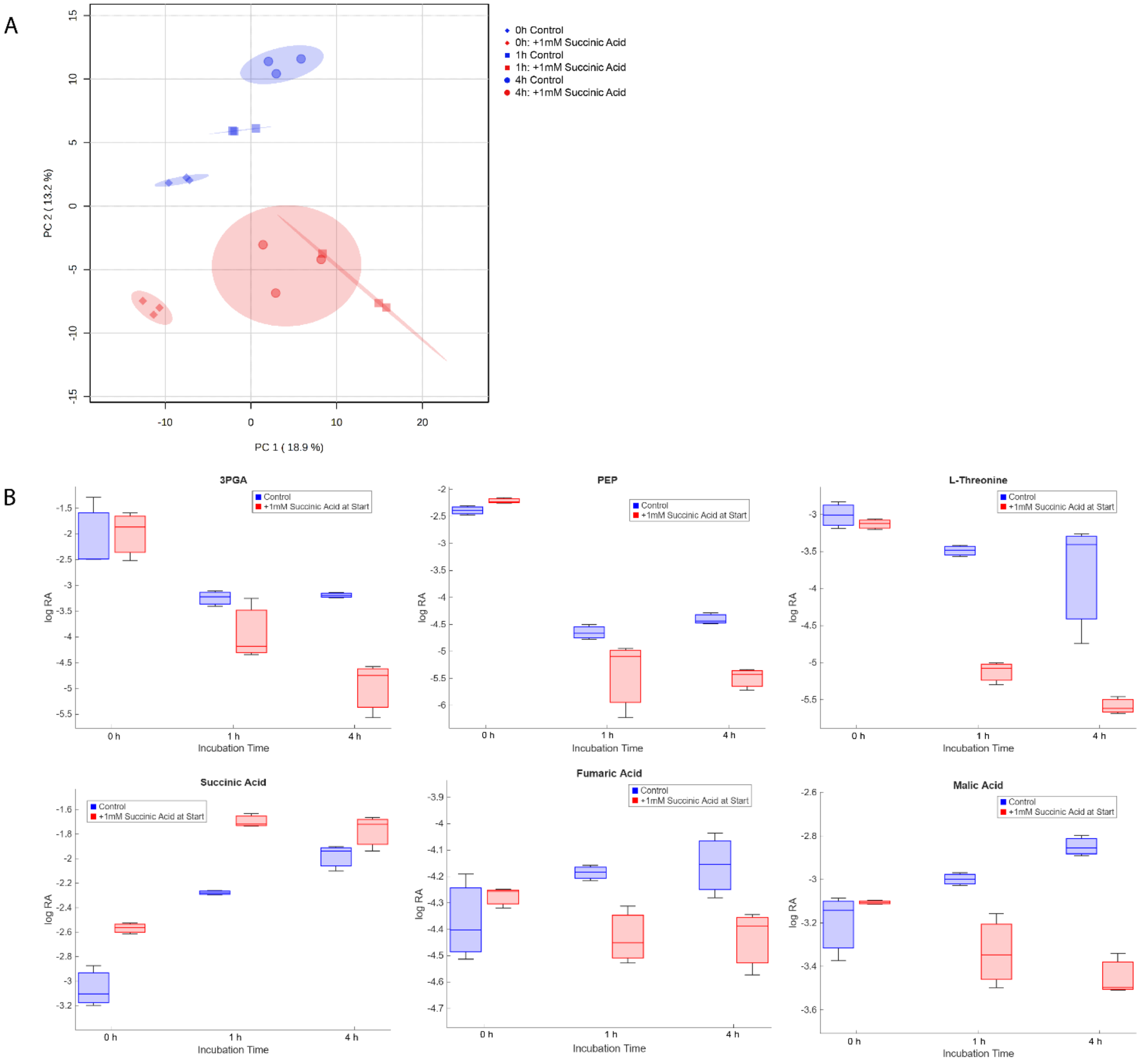


**Figure S7:** Metabolic effects of 1 mM succinic acid supplementation at 0 min. A) Multivariate analysis of the metabolome via PCA. Ellipses represent 95% confidence bounds. B) Some individual metabolites with different profiles between the supplemented and non-supplemented conditions. Blue bars represent medians, boxes represent 25$^{th}$ and 75$^{th}$ percentiles, whiskers indicate extreme values. While the dynamics of multiple other TCA cycle metabolites are substantially changed, so are some glycolytic metabolites and at least one amino acid. Of particular note is that succinic acid supplementation actually causes the qualitative profile of fumaric acid and malic acid to change, from accumulation in the control to depletion in the supplemented condition.

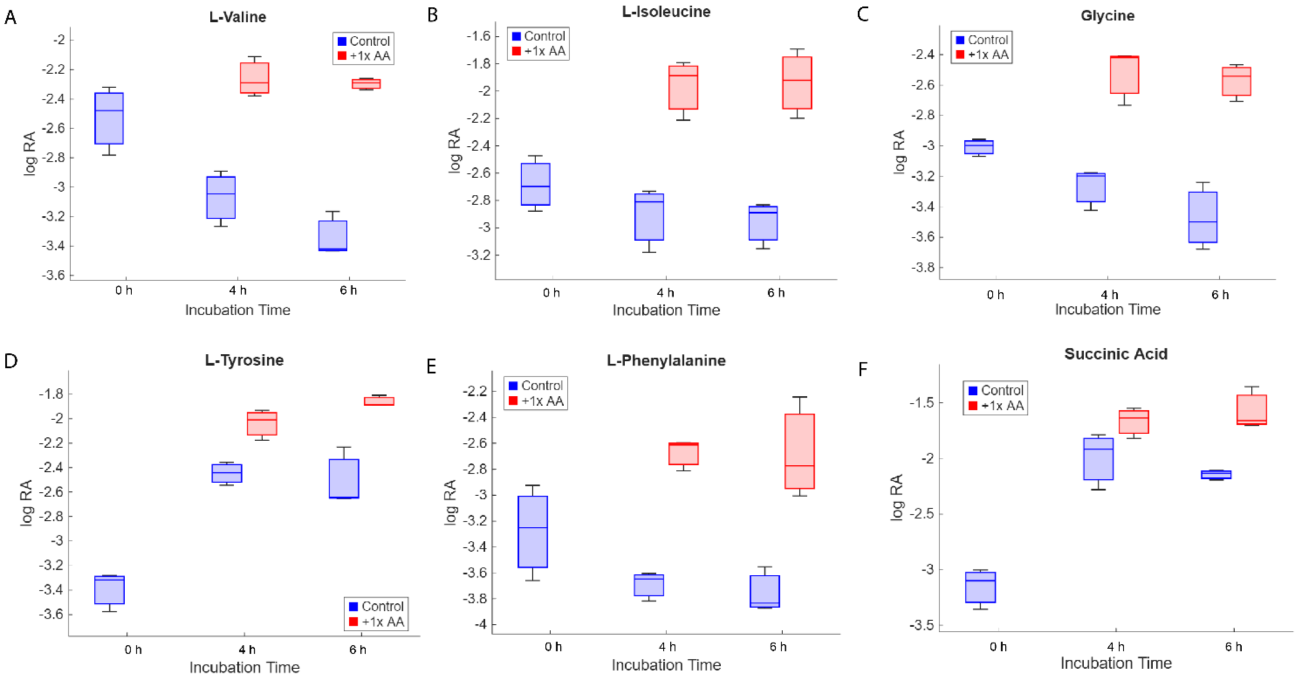


**Figure S8:** Additional individual metabolite results from Figure 4 for A) L-valine, B) L-isoleucine, C) glycine, D) L-tyrosine, E) L-phenylalanine, F) succinic acid. Blue and red bars represent medians for control and supplemented conditions, respectively, boxes represent 25th and 75th percentiles, whiskers indicate extreme values.

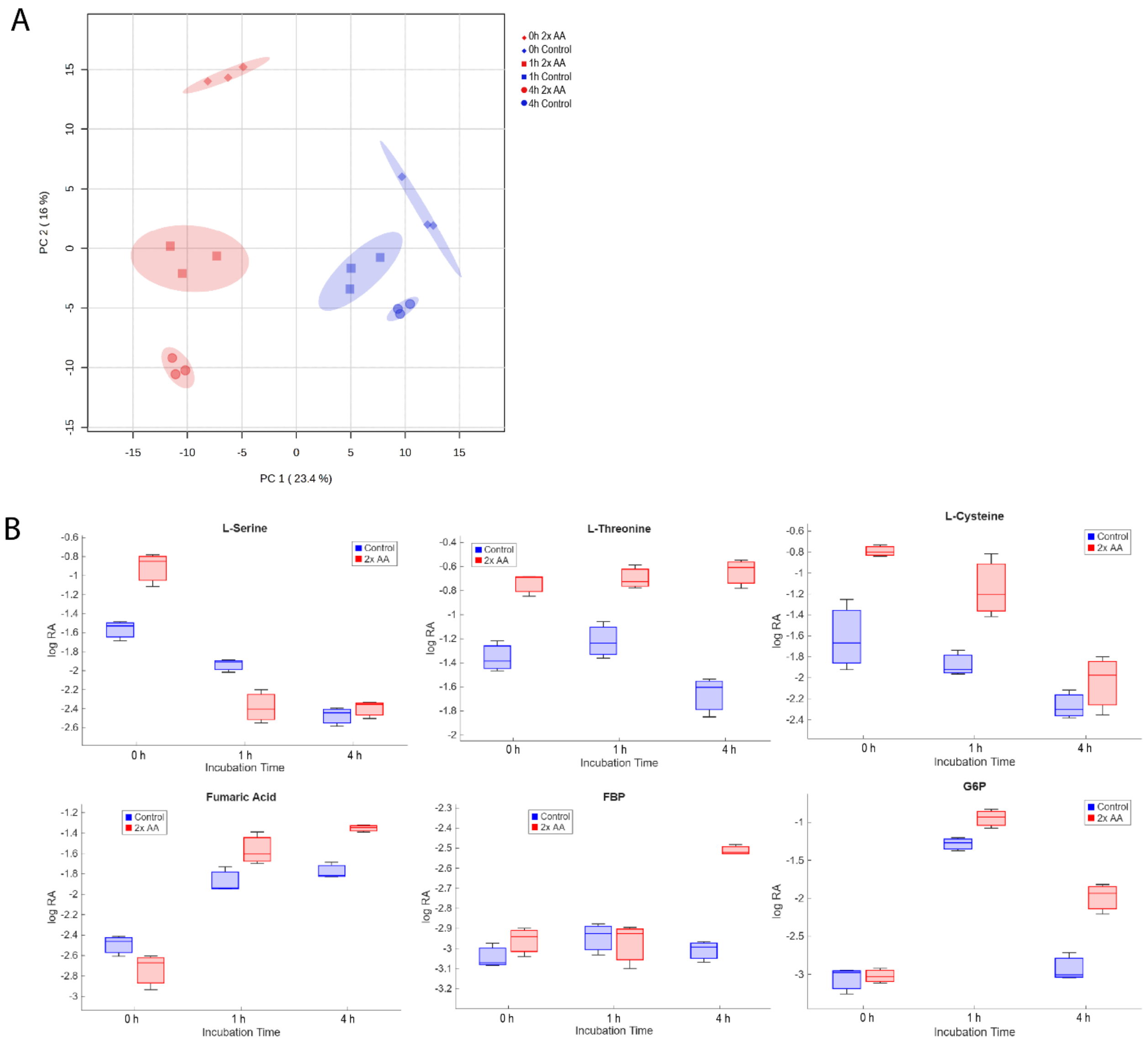


**Figure S9:** Metabolic effects of supplementing additional amino acids at 0 min. A) Multivariate analysis of the metabolome via PCA. Ellipses represent 95% confidence bounds. B) Some individual metabolites with different profiles between the supplemented and non-supplemented conditions. Blue bars represent medians, boxes represent $25^{th}$ and $75^{th}$ percentiles, whiskers indicate extreme values. Notably, unlike in Figure 4, the initial doubling of amino acids exaggerates systems-scale metabolic changes rather than decreasing them. Multiple metabolites appear to deplete faster despite starting at higher initial levels.
